# Finding the difference between periosteal and endocortical bone adaptation by using Artificial Neural Networks

**DOI:** 10.1101/357871

**Authors:** Abhishek Kumar Tiwari, Jitendra Prasad

## Abstract

In silico models of bone adaptation successfully simulated in vivo periosteal bone apposition, however, there are instances where these models may have limited success in predicting the new bone formation at endocortical surface. In vivo studies have highlighted that cortical bone surfaces may have differences in their modeling or remodeling responses to mechanical loading. However, the principle which the two cortical surfaces follow in bone adaptation is not very clear. This work accordingly attempts to understand how periosteal and endocortical surfaces accommodate loading-induced new bone formation. A neural network model is used to serve the purpose. A relationship is established to compute new bone thickness as a function of mechanical parameters (normal and shear strains) and non-mechanical parameters (distances from the neutral axis and the centroid) at the two surfaces. Analytical results indicate that two cortical surfaces behave opposite to each other in order to achieve optimal distribution of newly formed bone. The outcomes may be useful in establishing a unifying principle to predict site-specific new bone formation.

## 1. Introduction

Low-amplitude and cyclic mechanical loading may effectively inhibit or cure bone loss [1,2], as elevated normal strain magnitude promotes osteogenesis i.e. new bone formation [3]. Accordingly, in silico studies on cortical bone adaptation have assumed normal strain or strain energy density (SED) as the prime mechanical stimulus to model loading-induced osteogenesis [4–7]. These models have closely predicted the sites of new bone formation at periosteal surface. However, there are instances where computer models may have limited success in fitting new bone formation, especially at the endocortical surface. For example, Srinivasan et al. [8] accurately simulated bone formation at periosteal surface of a murine tibia mid-diaphyseal cross-section; however, the model has not correctly predicted osteogenesis at endocortical surface. They have concluded that cells involved in adaptation at endocortical surface may be different from periosteal precursors. Thus, the distinct milieu of these cells at the two cortical surfaces may have resulted in different new bone response. In addition, endocortical surface is typically subjected to marrow pressurization, which may have indeed contributed into a different response. Srinivasan et al. [8] also suggested that a planned computer model may be required to simulate osteogenesis at endocortical surface. Tiwari and Prasad [9] also modeled osteogenesis for an *in vivo* cantilever loading study on bone adaptation. The model successfully predicted site-specific new bone formation at the periosteal surface when normal strain is considered as the stimulus. The model, however, did not fit osteogenesis noticed at the endcortical surface especially near the neutral axis of bending; whereas, normal and shear strains in combination predicted site-specific new bone formation close to experimental values. This evidently indicates that the two cortical surfaces adapt differently to the same stimulus.

Judex et al. [10] reported that loading induces bone apposition at those sites where mechanical strength is least challenged such as at the neutral axis of bending. The possible reason may be the bone addition at such locations would preserve already-established physiological strain environment against any sudden change in loading pattern. Recent in vivo studies by Birkhold et al. [11,12] also substantiated that periosteal and endocortical bone surfaces exhibit distinct new bone response. Endocortical surface has been found more mechano-responsive to osteogenesis than the periosteal surface, and this higher endocortical mechano-responsiveness was independent of normal strain magnitude. It is evident from these studies that new bone formation may depend on non-mechanical factors such as spatial locations which may be due to regional differences in their site-specific biological/mechanical environment. However, the principle which decides the site-specificity of new bone formation at cortical bone surfaces is not well known. Accordingly, this work mainly aims to understand how differently two cortical surfaces behave with respect to bone adaptation. To understand this, it is also required to explore how cortical surfaces respond to mechanical and non-mechanical factors. These questions have been addressed in the present study using a neural network model.

In recent years, neural network model emerged as an efficient tool to establish the unforeseen relationships. There are in silico studies which have used neural network models in combination with finite element method to establish the relationship between bone modeling and mechanical environment. For example, Hambli [13] used finite element method and artificial neural network model together to predict adaptation in trabecular bone density. Barkaoui et al. [14] have also simulated the change in elastic material properties of trabecular bone using the similar approaches. In addition, Zadpoor et al. [15] used neural network model for predicting trabecular bone morphology in response to mechanical signals. These in silico studies have successfully simulated modeling and remodeling activities in trabecular bone. Our interest, however, is on cortical bone adaptation (in contrast to trabecular one) and only two studies have used neural network approach till present to analyze cortical bone adaptation [16,17]. A similar neutral network method has been used in the present study, however with different input parameters. The mechanical input parameters (i.e. normal and shear strains) are kept the same as that in our previous study [9], where there was a difference found in periosteal and endocortical bone adaptation. The present study attempts to investigate that difference.

We used the neural network approach to establish how cortical bone adaptation relates to mechanical and spatial (non-mechanical) parameters. The new bone thickness has been taken as the measure of cortical bone adaptation. Normal and shear strains are considered as mechanical parameters. Distances from the neutral axis and the centroid are kept as spatial parameters. A back-propagation neural network model is used with these parameters along with an assumption that periosteal and endocortical surfaces opt different mechanisms in deciding the sites and the amount of new bone formation. The model is trained with experimental data extracted from an in vivo study where tibia of a female C57Bl/6J mouse (10 weeks old) was subjected to cantilever cyclic loading of 0.5 N (‘high magnitude’) for 500 cycles at 1 Hz [18]. The model fits the extracted experimental data (i.e. new bone thickness, the site, as well as the amount and form of the mechanical environment) in an equation. This equation calculates the site-specific new bone thickness as a function of mechanical and spatial parameters close to experimental values of other cantilever loading studies. Therefore, the relationship can also be further analyzed to understand how site-specificity and the amount of new bone formation are regulated at the two cortical surfaces. The analysis shows that spatial fields such as distance from the neutral axis and distance from the centroid also decide new bone response, and this response may change with respect to the two cortical bone surfaces. This analysis also indicated that new bone formation is site-specifically regulated rather than only by mechanical environment. The findings may be beneficial in the development of robust computer model to predict loading-induced osteogenesis which may be further useful in design of effective biomechanical strategies to add new bone mass at a desired location e.g. the site where the bone loss occurred.

## 2. Methods

### 2.1. Problem definition

We intend to find out how periosteal and endocortical surfaces adapt differently to mechanical loading. The steps are as follows:

- Build a neural network model to predict site-specific new bone thickness as a function of mechanical and spatial parameters. This will include experimental data collection, designing of the network and preparation of the input parameters.
- Test this model by predicting the site-specific osteogenesis for other in vivo studies.
- Plot and analyze new bone thickness at periosteal vs. endocortical surfaces with respect to mechanical and spatial parameters in order to understand if the two surfaces behave in a similar or different manner.

### 2.2. Experimental data collection

We trained the neural network with the experimental data obtained from an in vivo study of Srinivasan et al. [18] where a murine tibia of C57Bl/6J female mouse (10 weeks old) was subjected to cyclic cantilever bending, and the new bone formation has been quantified at a mid-diaphyseal cross-section (Fig 1A). The cross section’s geometry with labeled new bone thickness is obtained from the histological images reported in the literature. The cross section under consideration has been divided into 36 equal-angle pie sectors which intersect periosteal and endocortical surfaces at 36 circumferential points (Fig 1B). Some of these data sets have been used for training, while the remaining data are verified post-training. These circumferential points serve as 36 data sets where each data set consists of input and output parameters.

**Fig 1.**
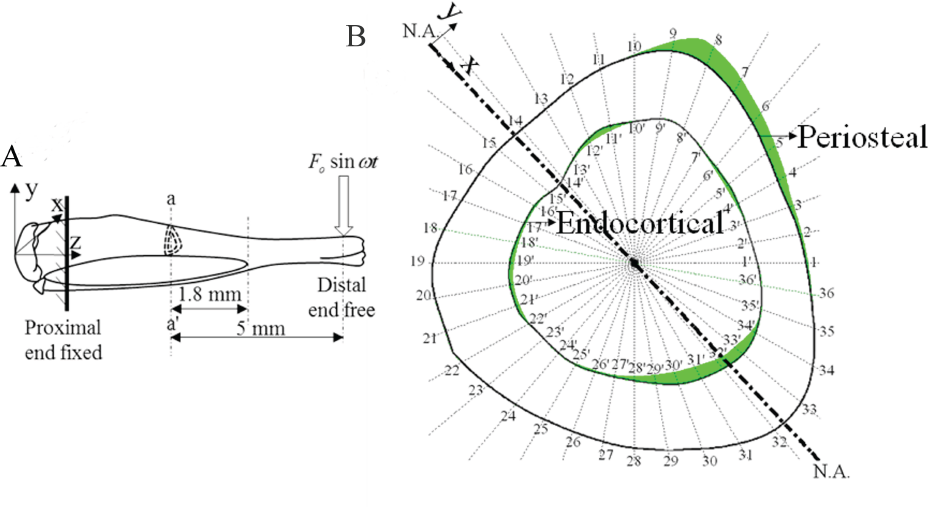
(A) In vivo configuration of murine tibia, and (B) new bone distribution (in green) at mid-diaphyseal cross section a-a’ (adapted from Srinivasan *et al.* [18]).

### 2.3. The network

A back propagation neural network approach with the single hidden layer is used to relate the site-specific new bone thickness to the corresponding spatial and mechanical parameters. Sigmoid transfer function has been selected as the activation function for the hidden layer since characteristics of this function closely represent the evolution law of bone surface adaptation [5]. Two spatial parameters (i.e. distances from the neutral axis and that from the centroid) along with two mechanical parameters (i.e. normal and shear strains) have been incorporated as a total of four input parameters to the model, whereas the corresponding new bone thickness has been taken as an output parameter. Thus, the network has 6 nodes in the input layer and one node in the output layer (Fig 2). MATLAB (Mathworks Inc. Boston) is used to program the neural network model.

**Fig 2.**
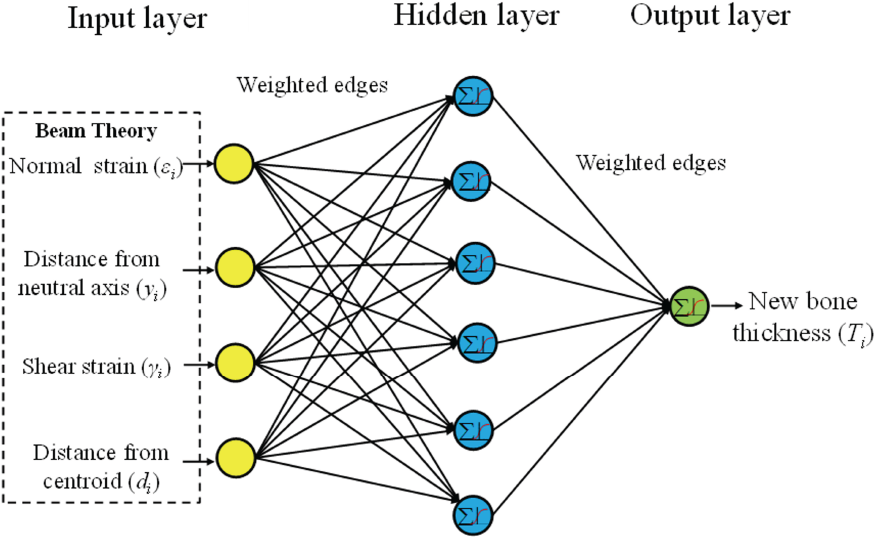
The neural network model to predict site-specific new bone thickness.

### 2.4. Preparation of input parameters

The distance from the neutral axis and that from the centroid were computed using standard coordinate geometry formulae. Normal and shear strains were estimated using the beam theory. Following relations have been used to compute normal and shear strains:

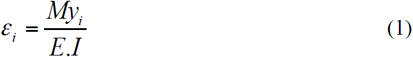

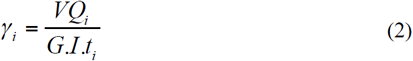

where, *ε*_*i*_ and *γ*_*i*_ are normal and shear strains, respectively, at any circumferential point i; *M* is bending moment at the cross section; *y*_*i*_ is the distance of any circumferential point from the neutral axis; *I* is the second moment of area; *V* is the shear force; *Qi* is the first moment of the area lying above an axis parallel to the neutral axis and passing through point i; *t_i_* is the total width of the section above afore-mentioned axis; and, *E* (20 GPa) and G (7.69 GPa) are Young’s modulus and shear modulus, respectively. Beam theory calculations have shown a maximum normal strain magnitude of approximately 1311 µε which was found close to in vivo strain values i.e. 1330 ± 50 µε [9].

### 2.5. Training and testing

The network has been trained using 24 data sets. Rest 12 data sets have been used for testing. Based on the assumption that cortical bone surfaces behave differentially to each other, the training has been individually performed for both the surfaces. The model has been further tested to predict new bone thickness noticed in three other cantilever loading studies [8,19]. Loading protocols of simulated studies are listed in Table 1. Based on loading protocols and histological cross sections reported in these studies, input parameters have been prepared and the new bone thickness has been calculated at periosteal and endocortical surface points.

**Table 1.** Loading parameters of simulated in vivo studies

| Protocol | Load (N) | Frequency (Hz) | Cycles | In vivo Study |
| --- | --- | --- | --- | --- |
| 1 | 0.5 | 1 | 450 | Srinivasan *et al.* [8] |
| 2 | 0.5 | 1 | 450 | Srinivasan *et al.* [8] |
| 3 | 0.3 | 1 | 1000 | Hankenson *et al.* [19] |

### 2.6. Analysis of the relationship

Training determines an optimal empirical relation of the new bone thickness to all the four input parameters. The relation obtained has been tested further for accuracy and analyzed as well. Three input parameters were fixed at one of the three sets of values according to Table 2, while remaining one parameter is varied and corresponding output (new bone thickness) has been plotted and analyzed. The response has also been compared with the experimental findings available in the literature.

**Table 2.** Fixed values of the parameters

| Parameters | Periosteal surface |  |  | Endocortical surface |  |  |
| --- | --- | --- | --- | --- | --- | --- |
|  | Max. | Mid. | Min. | Max. | Mid. | Min. |
| Distance from the neutral axis, $y$, mm | 0.6 | 0.35 | 0.1 | 0.5 | 0.25 | 0.05 |
| Normal strain, $\epsilon$, $\mu\epsilon$ | 1300 | 650 | 100 | 1000 | 500 | 50 |
| Shear strain, $\gamma$, $\mu\epsilon$ | 255 | 135 | 20 | 250 | 135 | 20 |
| Distance from the centroid, $d$, mm | 0.7 | 0.5 | 0.3 | 0.6 | 0.4 | 0.2 |

## 3. Results

### 3.1. Training

The neural network model establishes an empirical relationship for the site-specific new bone thickness (*T_i_*) as a function of spatial and mechanical parameters (i.e. distances from the neutral axis (*y_i_*) and the centroid (*d_i_*), normal strain (*ε_i_*), and shear strain (*γ_i_*)) which is as follows:

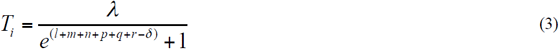

where

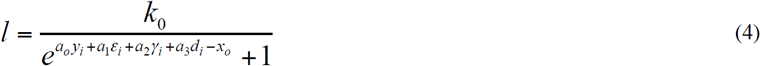

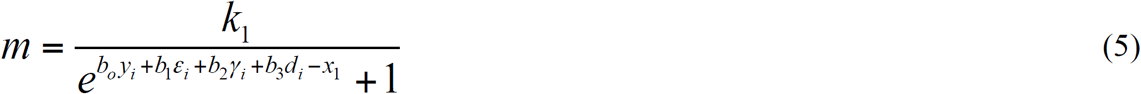

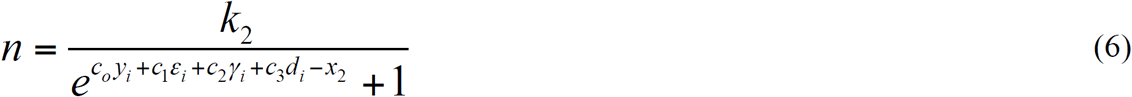

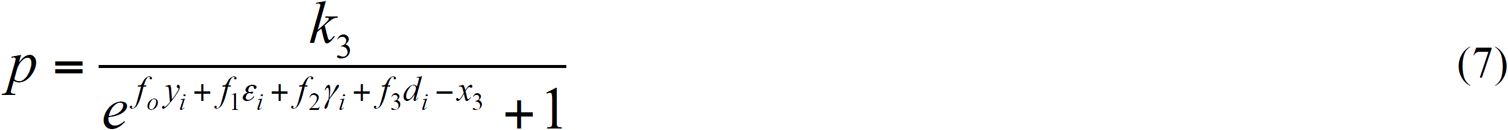

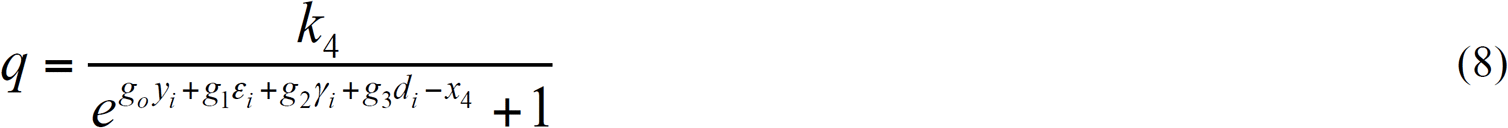

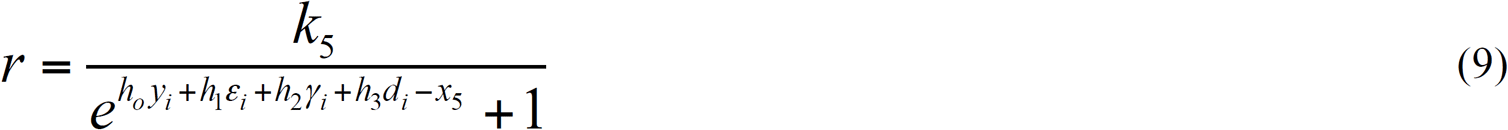

*a_n_, b_n_, c_n_, f_n_, g_n_* and *h_n_* (*n* = 0to3)*k_m_ and x_m_* (*m* = 0 to 5) are constants. The values of constants are different for periosteal and endocortical surfaces as given in Table 3.

### 3.2. Validation

The equation obtained from training is used to calculate new bone thickness for rest 12 circumferential points at periosteal and endocortical surfaces. Computed new bone thickness has been found close to experimental thickness noticed in the in vivo study [18]. The mean square error between computational and experimental results is nearly 6% (Fig 3). Fig 4 presents computational predictions of new bone distribution for all the three in vivo studies listed in Table 1. Computational new bone distribution is site-specifically compared with experimental distribution at 36 circumferential points. Fig 4A indicates the computational prediction of new bone thickness for “Protocol 1”. Computational new bone formation at periosteal points 1-7 and 18-23, and endocortical points 18′-36′ are found in accordance with experimental results. The relationship, however, overestimates osteogenesis at endocortical points 9′-18′. Similarly for “Protocol 2”, the new bone thickness estimated at periosteal points 1-12, 15-29 and 34-36, and endocortical points 6′-12′, 16′-22′ and 24′-36′ also align with the sites of in vivo new bone formation (Fig 4B). However, a slight overestimation is noticed at endocortical points 1′-6′, 16′-22′ and 24′-36′. The relationship also predicts new bone formation for “Protocol 3” at periosteal points 1-12 and 26-36, and endocortical points 4′-10′ (Fig 4C). Computational predictions are found close to experimental values of the new bone thickness except at periosteal points 12-24. It is, therefore, evident from results that the established relationship predicts in vivo new bone formation near and away from the neutral axis of bending or the sites of maximal and minimal normal strain magnitude. Thus, the incorporation of spatial parameters in the neural network model had successfully fitted the experimental data on loading-induced osteogenesis for those cases where other available bone adaptation models in the literature may have limitations.

**Fig 3.**
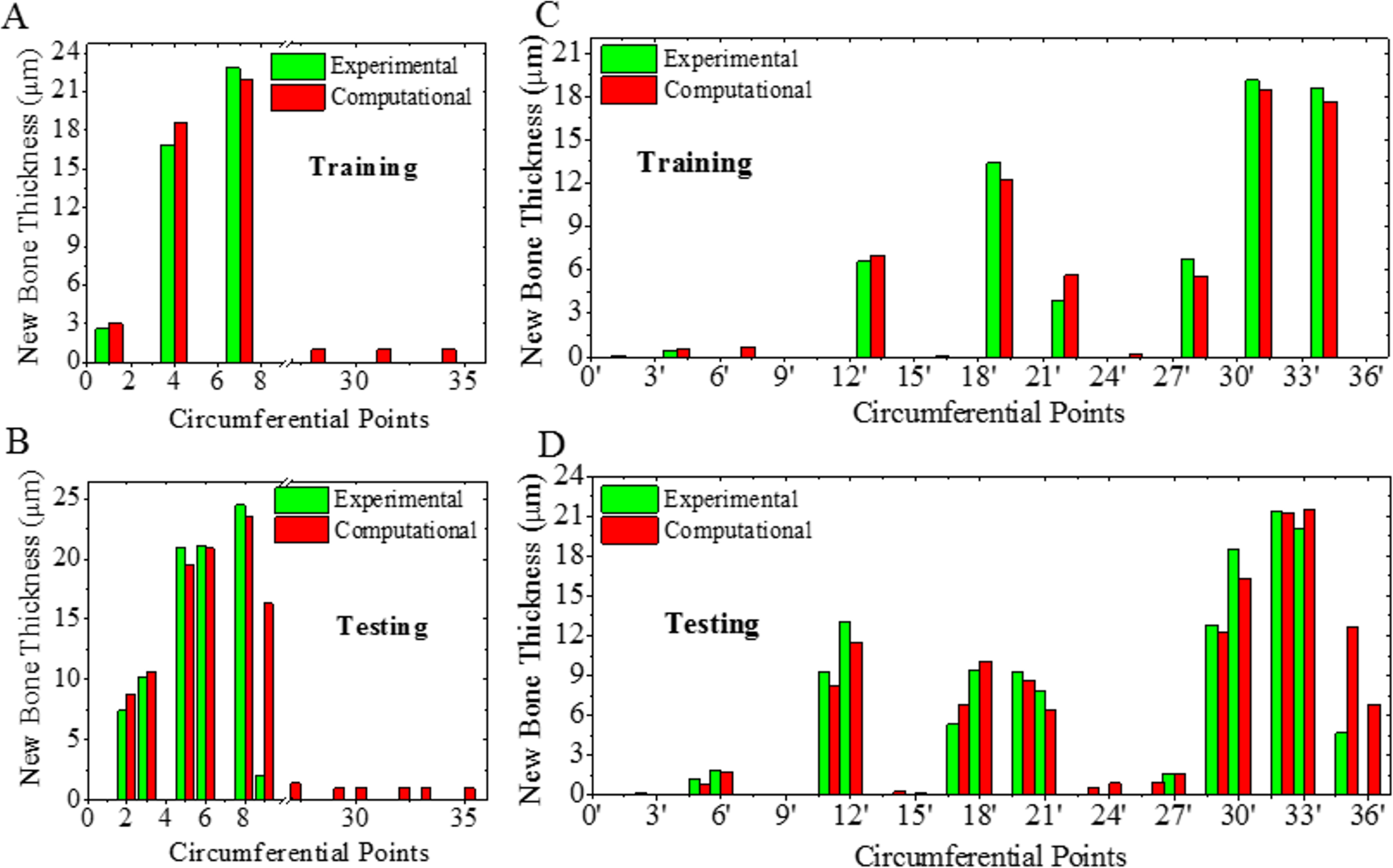
Site-specific experimental vs. computational new bone thickness in training and testing at: (A and B) periosteal and (C and D) endocortical surfaces. The new bone thickness absent on circumferential points is not shown.

**Fig 4.**
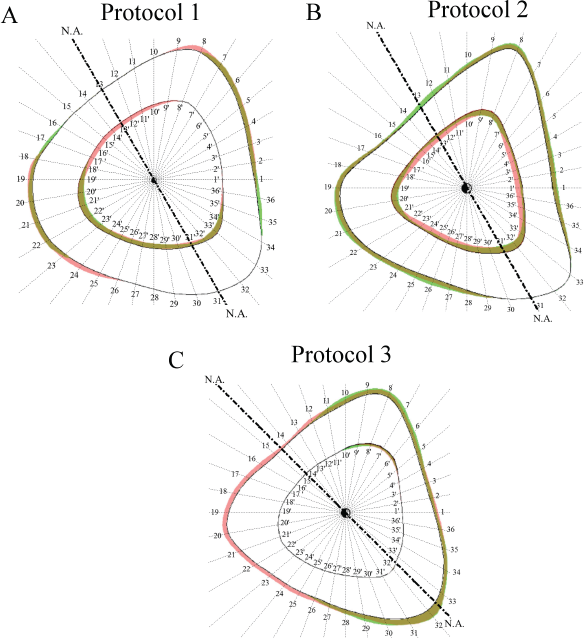
Computational (*red*) versus experimental (*green*) new bone distributions for (A) *Protocol 1*, (B) *Protocol 2* and (C) *Protocol 3* where in vivo new bone formation at murine tibiae midshaft cross sections are adapted from Figs 2(g) and 5(c) of Srinivasan *et al.* (2010) [8], and Fig 2(a) of *Hankenson et al.* (2006) [19]. The cross section’s circumference has been divided into 36 sectors in order to compare site-specific osteogenesis. The overlap of experimental and computational bone formation is presented in red-green mixture.

### 3.3. The analysis

This section presents the results obtained from the analysis. The results explain how new bone formation at periosteal and endocortical surfaces respond to spatial and mechanical parameters.

#### 3.3.1. Effect of input parameters at periosteal surface

i. *Distance from the neutral axis (y_p_)*: Figs 5A to 5C show the effect of distance from neutral axis on new bone thickness. It is apparent from the figures that the new bone thickness increases nonlinearly with an increase in the distance from the neutral axis and becomes approximately constant after a certain value of the distance. Similar trends have been noticed for any values of the other three fixed parameters, namely normal strain, shear strain and distance from the centroid. The negative values of *y_p_* i.e. the distance corresponding to the compressive side of bending, led to no new bone thickness. This indicates that most of the new bone formation coincides with tensile normal strain which has been noticed in the simulated in vivo studies. Birkhold *et al.* [11] have also noticed a greater cortical adaptive response in the regions experiencing tensile strain compared to compressive strains, which also aligns with the nature of established relationship.
ii. *Normal strain* (*ɛ_p_*): New bone thickness also increases nonlinearly with an increase in normal strain as well (Figs 5D to 5F). The trends in these figures are similar to that in Figs 5A to 5C, respectively, as expected from the beam theory. This characteristic also aligns with the observations of *Hseih and Turner* where a non-linear increase in bone formation rate is noticed with increase in load or strain magnitude [20].
iii. *Shear strain (γ_p_)*: As can be seen in Figs 5G to 5I, the new bone thickness decreases with increase in shear strain. In addition, maximum new bone thickness can be seen to occur at minimal shear strain (i.e., *γ_p_ = 0*), i.e. where the normal strain is maximal.
iv. *Distance from the centroid (d_p_)*: It can be observed in Figs 5J to 5L that new bone thickness increases nonlinearly with an increase in distance from centroid.

**Fig 5.**
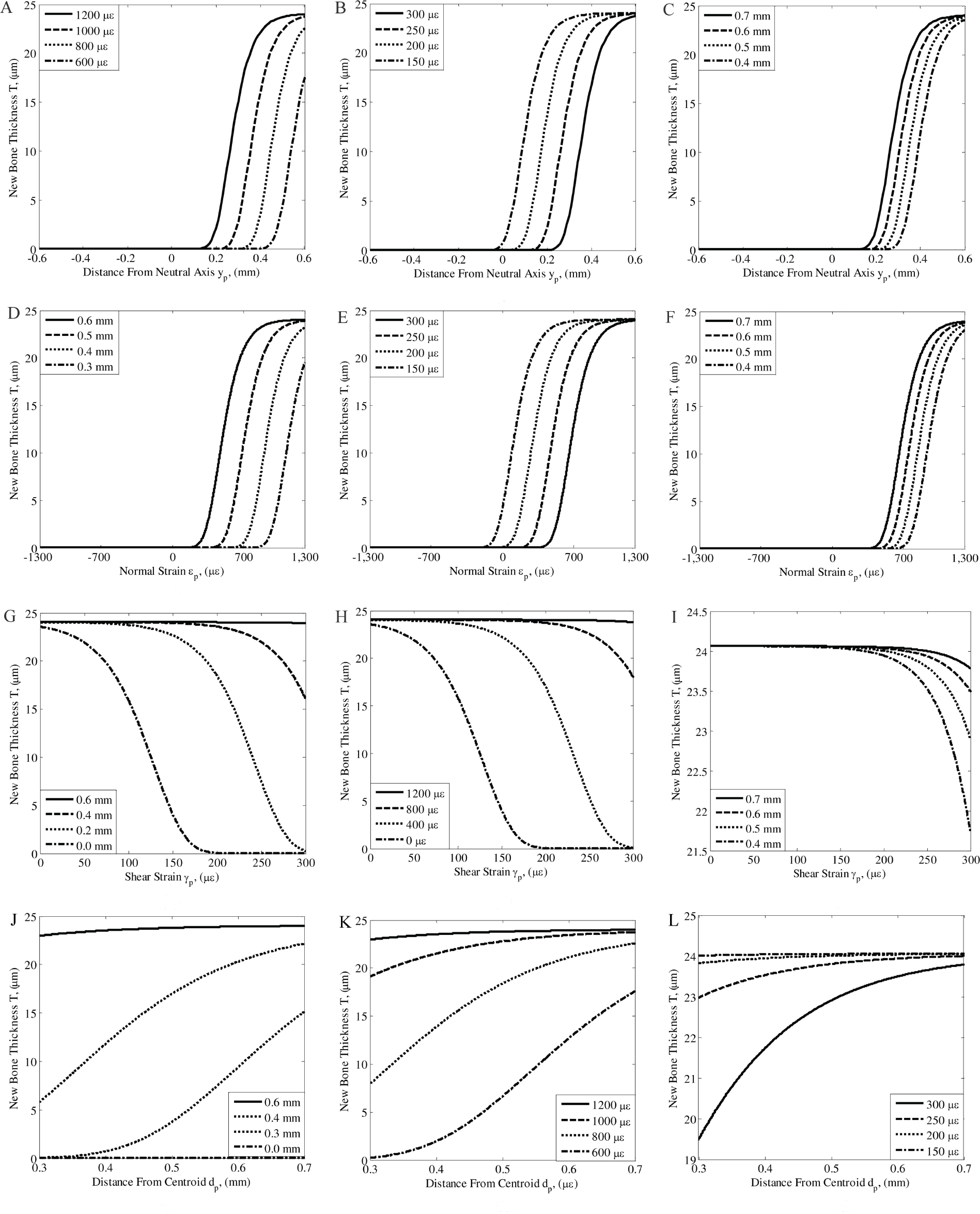
New bone thickness (*T*) vs. distance from the neutral axis (*y_p_*) at periosteal surface for different values of: (A) normal strain, (B) shear strain, and (C) distance from the centroid; *T* vs. normal strain (*ε_p_*) for different values of: (D) distance from the neutral axis, (E) shear strain, and (F) distance from the centroid; *T* vs. shear strain (*γ_p_*) for different values of: (G) distance from the neutral axis, (H) normal strain, and (I) distance from the centroid; and *T* vs. distance from the centroid (*d_p_*) for different values of: (J) distance from the neutral axis, (K) normal strain, and (L) shear strain.

#### 3.3.2. Effect of input parameters at endocortical surface

i. *Distance from the neutral axis (y_c_):* This is interesting to note that the effect of distance from the neutral axis on new bone formation at endocortical surface (Figs 6A to 6C) is mostly opposite of that at periosteal surface (Figs 5A to 5C). Unlike the periosteal surfaces, the new bone formation at endocortical surface is either at neutral axis or on the compressive side of the cortex.
ii. *Normal strain (ε_c_):* As normal strain is proportional to distance from the neutral axis, the effect of normal strain on new bone formation (Figs 6D to 6F) is the same as that of distance from the neutral axis (Figs 6A to 6C). Such responses align with findings of Birkhold *et al.* [11] where osteogenesis at endocortical surface occurred at a lower strain magnitude than the periosteal surface.
iii. *Shear strain (γ_c_):* The effect of shear strain on new bone thickness at endocortical surface (Figs 6G to 6I) is found to be roughly opposite of that at periosteal surface (Figs 5G to 5I). New bone thickness at endocortical surface increases nonlinearly with an increase in shear strain magnitude. Skedros *et al.* [21] also noticed a greater cortical thickness in the regions experiencing highest shear strain.
iv. *Distance from the centroid (d_c_):* It can be observed in Figs 6J to 6L that new bone thickness first increases with an increase in distance from the centroid, becomes maximum at a certain value, and then decreases onward back to zero. Larger is the normal strain, the nearer is the peak of bone thickness from centroid. In contrast, larger is the shear strain, the further is the peak of bone thickness from the centroid. Thus, normal strain or shear strain decides the optimal distance from centroid where new bone thickness would be the maximum.

**Fig 6.**
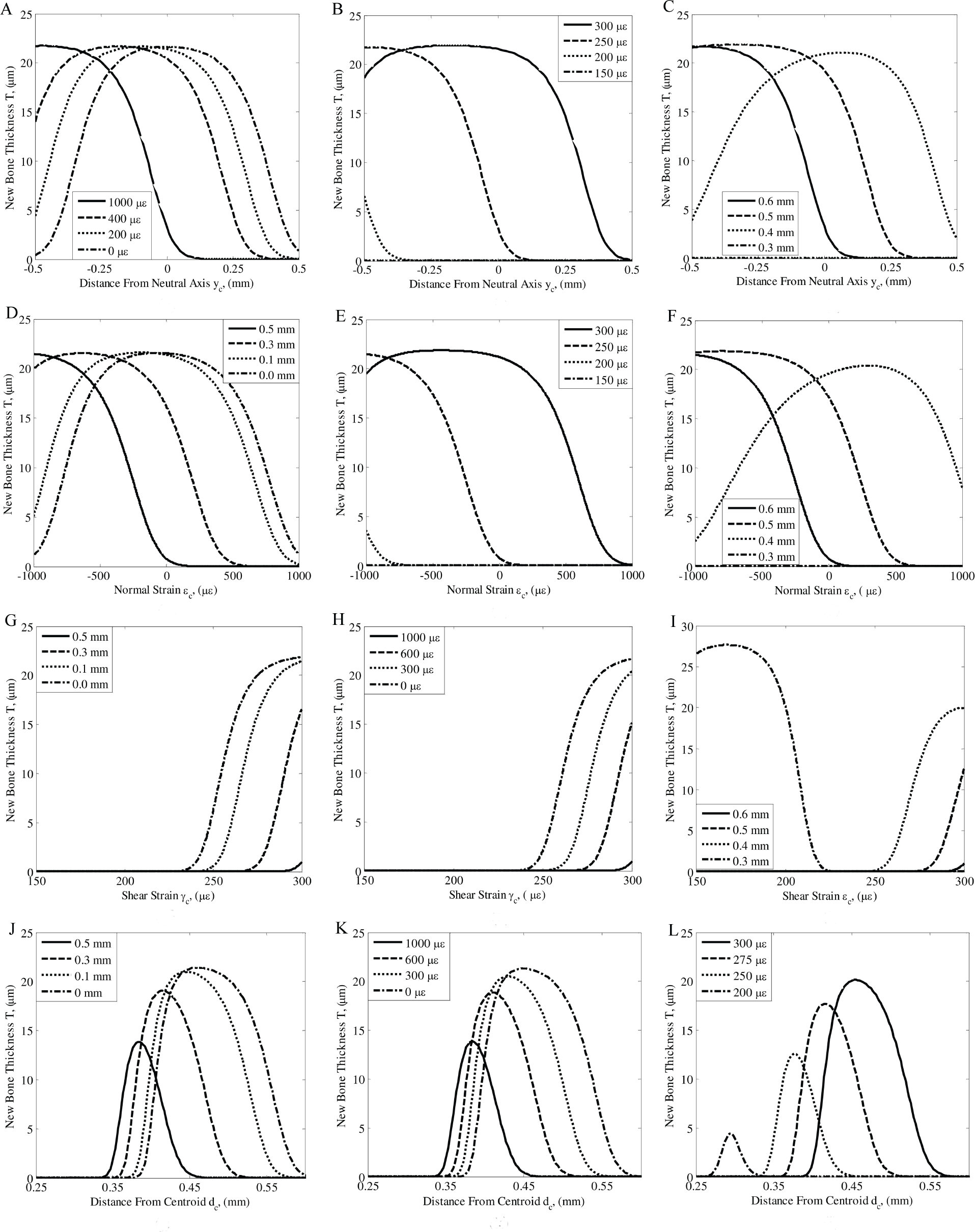
New bone thickness (*T*) vs. distance from the neutral axis (*y_c_*) at endocortical surface for different values of: (A) normal strain, (B) shear strain, and (C) distance from the centroid; *T* vs. normal strain (*ε_c_*) for different values of: (D) distance from the neutral axis, (E) shear strain, and (F) distance from the centroid; T vs. shear strain (*γ_c_*) for different values of: (G) distance from the neutral axis, (H) normal strain, and (I) distance from the centroid; and *T* vs. distance from the centroid (*d_c_*) for different values of: (J) distance from the neutral axis, (K) normal strain, and (L) shear strain.

In the above sub-sections, the response of new bone thickness at periosteal and endocortical surfaces has been analyzed when other three parameters were fixed to their maximum values. Hence, we have also studied the effect of each parameter by fixing other three parameters to their maximum, middle and minimum values, as given in Table 1. The new bone thickness at periosteal surface is largest for ‘Max’ set of values and smallest for ‘Min’ set of values (Figs 7A to 7D); however, the plots (Figs 7A to 7D) followed the similar trends as in Fig 5. Similarly, the effect of parameters on new bone thickness at endocortical surface (Figs 7E to 7H) is almost similar as in Fig. 6 and nearly opposite to that noticed at periosteal surface in Figs 7A to 7D, as thickness is found largest for ‘Min’ set of values.

**Fig 7.**
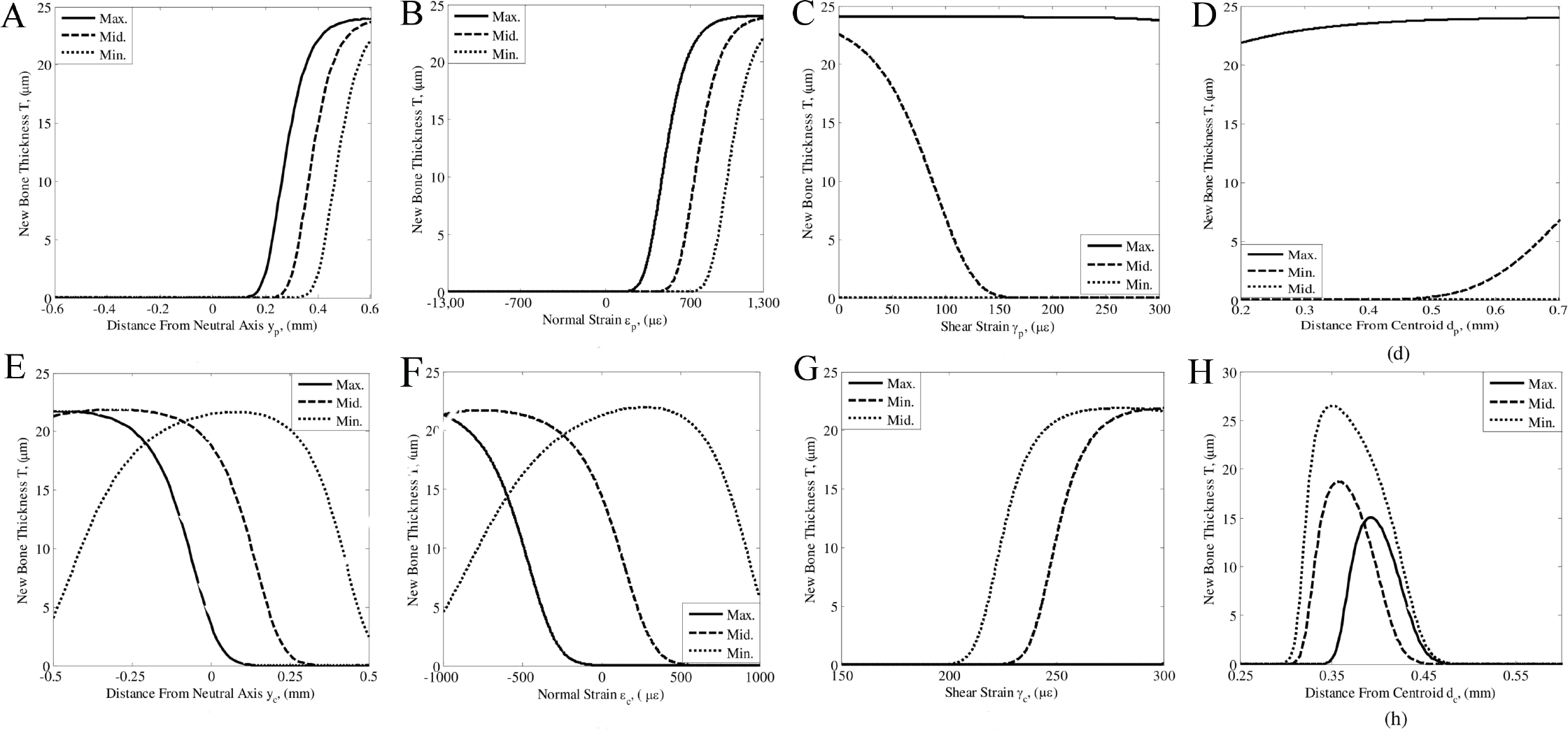
New bone thickness (*T*) vs. distance from the neutral axis (*y*), normal strain (*ε*), shear strain (*γ*) and distance from the centroid (*d*) for maximum, middle and minimum values of fixed parameters at (A, B, C and D) periosteal and (E, F, G and H) endocortical surfaces.

These plots particularly describe the adaptation characteristics of cortical bone surfaces. Periosteal surface allows new bone addition at the sites away from both of the neutral axis of bending and the centroid, or at the sites of maximal strain magnitude. On the other hand, endocortical surface allows osteogenesis at the sites near the neutral axis or at farthest endocortical sites from the newly formed bone at the periosteal surface. Thus, it is evident that periosteal surface closely follows Frost’s mechanostat theory of adaptation as new bone formation occurred above an osteogenic threshold in response to all the input parameters. However, it appears that endocortical surface does not follow this rule as it behaves opposite to the periosteal surface. These finding may be incorporated in a computer model of bone adaptation to precisely predict the loading induced osteogenesis.

## 4. Discussion

The model correctly estimates site-specific new bone thickness at cortical bone surfaces for in vivo cantilever loading studies when spatial and mechanical parameters have been used as inputs. The analysis suggests that new bone response at endocortical surface is typically opposite to that at periosteal surface. It has also been found that there is an optimal distance from the centroid for maximum new bone formation, where the optimal distance is jointly decided by normal and shear strains. These findings support available literature, as described below.

*Carpenter and Carter* [22] indicated that intracortical stresses govern bone size whereas periosteal surface pressure regulates sites of new bone formation. The periosteal loading environment/pressure is typically induced by structures surrounding the bone cross section, which is considered essential for the development of mid-diaphyseal cross section. Thus, surrounding structures may also be responsible for site-specific bone response, and bone also adapts to accommodate adjacent anatomical structures. It is therefore evident that bone response is not entirely governed by mechanical stresses or strains; the response may also vary site-specifically. Our findings also support this fact as the sites of osteogenesis at both the cortical surfaces (periosteal and endocortical) have shown dependency on spatial parameters. Previous studies have shown that bone apposition at periosteal surface always occurs along with bone resorption at the adjacent endocortical surface [23]. Hence, bone adaptation follows a specific modelling pattern i.e. formation and resorption at adjacent bone surfaces and vice-versa. Our analysis also substantiated this fact as the new bone responses to mechanical stimuli at endosteal surface have been found nearly opposite to that at the periosteal surface.

It is reported that periosteal surface pressure inhibits new bone formation required to align in the direction of maximal bending stiffness with applied bending moment direction [22]. The bone, therefore, adapts in such a way that minimum bending stiffness aligns with the direction of applied bending moment. Accordingly, new bone apposition intend to occur near the neutral axis or the sites of minimal normal strain magnitude to maintain minimal bending stiffness in the direction of habitual bending. This may be one of the possible reasons due to which endocortical surface attempts to accommodate maximum new bone thickness typically around the neutral axis of bending in contrast to the periosteal surface. *Lanyon and Baggot* [24] also noticed that longitudinal tensile strain and compressive strain may have opposite effects on bone remodeling. It indicates that periosteal surface pressure and intracortical stresses act collectively to deliver optimal new bone distribution over the cortex. Our analysis also suggests that maximal periosteal new bone formation occurs at tensile side, and the new bone thickness may also nonlinearly increase with increase in strain magnitude.

Fan *et al.* [25] have also indicated that physiological activities such as modeling and remodeling are optimization processes. They have concluded that bone’s adaptation is regulated by an optimal structure principle which depends on geometrical/spatial parameters such as centroid of the cross-section and center of mass [25,26]. In other words, bone adapts to exogenous loading by adjusting center of mass and centroid in an optimal way to enhance the load bearing capacity. Our study also suggests that most of new bone formation at the endocortical surface occurred within a specific range of distance from centroid. Hence, cortical bone cross-section has a tendency to either maintain optimal stiffness with respect to physiological loading or shift their area towards the centroid.

Skedros *et al.* [21] have noticed greater lacuna density and vascular porosity near the neutral axis of bending at endosteal surfaces. These sites experience elevated shear strain due to bending and a greater osteocyte population density was also found in this region. This indicates that maximum remodeling activities would occur in the region of minimal normal strain magnitude, and the same was noticed in terms of distribution of new bone thickness along the cortex. Thus, the cortical bone surface may also adapt differently to shear strain stimulus which has also been substantiated in this study.

There are very few studies in the literature, which have attempted to relate site-specific osteogenesis at cortical bone surfaces to mechanical environment using a neural network approach. For example, Mi *et al.* [16] have attempted to relate site-specific osteogenesis to strain gradients or fluid shear using network models, and accordingly a linear relationship is assumed in the study. However, their results indicated that new bone formation and mechanical parameters exhibit a non-linear relationship between them. A similar non-linear relationship has been exhibited by cortical bone surfaces which closely fitted the experimental data. Previous in silico studies on bone adaptation has considered only one mechanical stimulus; whereas, *Tiwari and Prasad* suggested that different mechanical stimuli (normal and shear strains or fluid shear) act collectively [9]. This work also substantiates that normal and shear strains along with spatial parameters are required to completely model the osteogenesis at the two cortical surfaces.

In addition to the exogenous mechanical environments, bone adaptation at the cortical surfaces is also governed by biological processes and local anatomical variables. Incorporation of spatial parameters (i.e. distance from the neutral axis and the centroid to the site of new bone formation) in the relationship serves to take care of such anatomical factors. The relationship is, however, derived and analyzed based on a limited experimental data, and new bone distribution has been computed for the cases of cantilever loading only. Bone remodeling data from diverse loading cases such as three point bending [27], four point bending [28], axial [29] and knee loading [30], will be incorporated in future to understand if the adaptation pattern of cortical surfaces changes with in vivo loading case. Loading parameters such as frequency, cycle, and rest insertion time, animal age, and other mechanical stimuli such as fluid shear, strain gradients etc. considerably influence the distribution of new bone thickness. The effects of these parameters have also not been studied in the context of adaptation at cortical surfaces. Nevertheless, there is hardly any study which addresses how the two cortical bone surfaces respond to loading-induced osteogenesis. This investigation demonstrates that periosteal and endocortical surfaces follow entirely opposite principles in accommodating the new bone as the periosteal surface was found more responsive to mechanical parameters while adaptation at the endocortical surface was governed by spatial parameters. Thus, it is recommended that different set of rules must be defined to predict site-specificity of new bone formation at cortical bone surfaces.

## 5. Conclusions

This work explains how periosteal and endocortical surfaces respond to mechanical environment which induces new bone formation. The present study establishes a relationship to estimate site-specific new bone thickness as a function of both spatial (distances from the centroid and the neutral axis) and mechanical parameters (normal and shear strains). The results indicate that new bone formation at cortical surfaces are regulated by mechanical as well as spatial parameters. This study concludes that periosteal and endocortical surfaces behave opposite to each other in order to achieve optimal distribution of new bone formation. The conclusions drawn here may be incorporated in the in silico models to precisely predict the site-specific new bone formation, which may be further utilized to design the prophylactic exercises to rescue bone loss or osteopenia.

## Acknowledgments

The authors acknowledge IIT Ropar for the institute fellowship. The authors acknowledge Mr. Dinesh Kumar and Mr. Santosh Kumar for their help in computation.

